# Amyloid-β_1–42_ oligomers enhance mGlu_5_R-dependent synaptic weakening via NMDAR activation and complement C3aR/C5aR signaling

**DOI:** 10.1101/2022.10.24.513427

**Authors:** Ai Na Ng, Eric W. Salter, John Georgiou, Zuner A. Bortolotto, Graham L. Collingridge

## Abstract

Synaptic dysfunction, weakening, and loss of synapses are well-correlated with the pathology of Alzheimer’s disease (AD). Oligomeric amyloid beta (oAβ) is considered a major synaptotoxic trigger for AD. Recent studies have implicated hyperactivation of the complement cascade as the driving force for loss of synapses caused by oAβ. However, the initial synaptic cues that trigger pathological complement activity remain elusive. Here, we examined a form of synaptic long-term depression (LTD) mediated by metabotropic glutamate receptors (mGluR) which is disrupted in rodent models of AD. Exogenous application of oAβ (1-42) to mouse hippocampal slices enhanced the magnitude of mGlu subtype 5 receptor (mGlu_5_R)-dependent LTD. We found that the enhanced synaptic weakening occurred via both NMDARs and complement C3aR/C5aR signaling. Our findings reveal a mechanistic interaction between mGlu_5_R, NMDARs, and the complement cascade in synaptic weakening induced by oAβ, which could represent an early trigger of synaptic loss and degeneration in AD.

## Introduction

Alzheimer’s disease (AD) is a progressive neurodegenerative disease characterized by memory loss, cognitive deficits and changes to personality and behavior (Breijyeh & Karaman, 2020). Although it is the most common cause of dementia affecting the aging population worldwide, therapeutic options are limited. Weakening and subsequent loss of synapses are early events in AD progression that precede the deterioration of memory and cognitive functions (Selkoe, 2002; Koffie, Hyman, & Spires-Jones, 2011; Li & Selkoe, 2020). The development of improved treatment options requires a deeper understanding of how synapses are initially impaired and how this leads to overt neuronal degeneration.

A key pathological feature of AD are plaques, which are made up primarily of insoluble aggregates of amyloid beta (Aβ) protein derived from the amyloid precursor protein (APP). The original amyloid cascade hypothesis posited that these Aβ plaques are the initial trigger of AD pathology (Hardy & Higgins, 1992). However, subsequent research found that the amount of soluble oligomers of Aβ (oAβ) better correlates with the degree of cognitive impairment in AD patients (DeKosky & Scheff, 1990; Terry et al., 1991). Interestingly, oAβ binds to excitatory synapses (Lacor et al., 2004; Lacor et al., 2007; Koffie et al., 2009), leading to disruption of synaptic function. Normally, synaptic strength is modulated through long-term potentiation (LTP) and long-term depression (LTD), which are thought to provide the cellular and molecular basis of learning and memory (Bliss & Collingridge, 1993). oAβ mediates a shift in the LTP/LTD balance, favoring LTD over LTP, leading to eventual net synapse loss (Lambert et al., 1998; Kim, Anwyl, Suh, Djamgoz, & Rowan, 2001; Hsieh et al., 2006; Shankar et al., 2007; Shankar et al., 2008; Li et al., 2009; Jo et al., 2011; Cline, Bicca, Viola, & Klein, 2018; Li & Selkoe, 2020; Taylor, Emptage, & Jeans, 2021). Therefore, the study of oAβ-dependent changes to LTD provides a means to investigate the initial synaptic changes relevant to the earliest stages of AD.

Synapse damage and loss caused by oAβ does not occur in a neuron-autonomous manner. The complement cascade is an innate immune pathway which has emerged as a central driver of synapse loss in AD (Wu, Dissing-Olesen, MacVicar, & Stevens, 2015; Hammond, Robinton, & Stevens, 2018). Complement cascade activation leads to phagocytosis by microglia via C3b/CR3, as well as chemotactic and inflammatory signaling via C3a-C3aR and C5a-C5aR axes (Ricklin, Hajishengallis, Yang, & Lambris, 2010). Tissue from post-mortem samples of patients with AD demonstrates activation of complement activity, including increased deposition of C1q and proteolytic cleavage of C3 (Afagh, Cummings, Cribbs, Cotman, & Tenner, 1996; Yasojima, Schwab, McGeer, & McGeer, 1999; Wu et al., 2019). Importantly, oAβ acts as a potent activator of the complement cascade and oAβ-mediated synapse loss is dependent on the complement pathway (Hong et al., 2016; Bie, Wu, Foss, & Naguib, 2019). However, the upstream synaptic signals that lead to aberrant activation of complement and subsequent synapse loss induced by oAβ are not well understood.

LTP and LTD of glutamatergic synapses is mediated by activation of N-methyl-D-aspartate receptors (NMDARs) and/or metabotropic glutamate receptors (mGluRs: Collingridge, Peineau, Howland, & Wang, 2010; Nicoletti et al., 2011; Volianskis et al., 2015). LTD, induced either by NMDARs or mGluRs, can trigger subsequent synapse elimination (Shinoda, Tanaka, Tominaga-Yoshino, & Ogura, 2010; Wiegert & Oertner, 2013; Hasegawa, Sakuragi, Tominaga-Yoshino, & Ogura, 2015; Wiegert, Pulin, Gee, & Oertner, 2018). Both mGluR-and NMDAR-dependent signaling are affected by oAβ, leading to synaptic plasticity impairments and aberrant synapse loss (Hsieh et al., 2006; Shankar et al., 2007; Shankar et al., 2008; Jo et al., 2011; Cavallucci et al., 2013; Beraldo et al., 2016; Jackson et al., 2019; Abd-Elrahman et al., 2020; Li & Selkoe, 2020).

Synaptic plasticity involving the mGluR subtype 5 (mGlu_5_R) is particularly pertinent to the understanding of AD for multiple reasons. For example, mGlu_5_R was been found to be the only co-receptor of cellular prion protein (PrP_c_) necessary for oAβ to activate intracellular signaling in neurons (Um et al., 2013). Further, both knockout of mGlu_5_R and pharmcological antagonism is protective in oAβ-based and genetic models of AD (Hamilton, Esseltine, DeVries, Cregan, & Ferguson, 2014; Hu et al., 2014; Hamilton et al., 2016; Abd-Elrahman et al., 2020). Interestingly, the complement cascade has recently been implicated in oAβ-dependent synapse loss mediated by mGluR activation (Bie et al., 2019). A silent allosteric modulator (SAM) of mGlu_5_R has also been found to restore synapse density through reduced tagging by C1q and microglia engulfment in a genetic mouse model of AD (Spurrier et al., 2022).

Previously, we identified a simple method to study mGlu_5_R-mediated LTD, via the brief application, S-3,5-dihydroxyphenylglycine (DHPG) which causes lasting depression of AMPAR-mediated synaptic transmission (Palmer, Irving, Seabrook, Jane, & Collingridge, 1997). This DHPG-induced LTD (DHPG-LTD) has since been used extensively to uncover many aspects of mGluR physiological and pathological function, including the regulation of mGluR function in AD mouse models (Hsieh et al., 2006; Shankar et al., 2008; Li et al., 2009; Cavallucci et al., 2013; Mango & Nisticò, 2018; Yang, Zhou, Ryazanov, & Ma, 2021). Here, we have used DHPG-LTD to investigate how the complement cascade may be rapidly recruited by mGluR activation. We have applied oAβ as a standard way to induce synaptotoxicity that is relevant to AD pathology. We found that oAβ enhanced mGlu_5_R-mediated LTD via a mechanism that involves both the activation of NMDARs and the C3aR and C5aR arms of the complement cascade. These findings reveal a signaling axis between glutamate receptors and the complement cascade which triggers early synaptic dysfunction that may underlie the initial stages of dementia.

## Materials and Methods

### Animals

Adult C57BL/6JOlaHsd male mice (10-16 weeks old) from Envigo (Bicester, U.K.) and a mGlu_5_R knockout line (Lu et al., 1997), backcrossed more than 10 generations to a C57BL/6J background, were used in this study. We chose to use male mice due to the recently reported sex specificity of oAβ-mGlu_5_R binding which is present only in males (Abd-Elrahman et al., 2020). Animals were housed four per cage in a room with designated 12 h light/dark cycle (lights on at 08:15/ off at 20:15) with temperature at 21 °C ± 2 °C and access to food and water *ad libitum.* All procedures were performed in accordance with the U.K. Animals (Scientific Procedures) Act of 1986. Genomic DNA from mGlu_5_R knockout litters were isolated from tail tips with DNeasy blood and tissue kit (Qiagen; Hilden, Germany; cat. no. 69504) and amplified as described (Lu et al., 1997). The amplified products were resolved and visualized in ethidium bromide-stained agarose gel.

### Oligomeric amyloid-β_1-42_ (oAβ) preparation

oAβ was prepared as described (Whitcomb et al., 2015). Briefly, Aβ peptide_(1-42)_ (MilliporeSigma; Burlington, Massachusetts; TFA recombinant human, ultra-pure; cat. no. AG912) was dissolved in 100 % 1,1,1,3,3,3-hexafluoro-2-propanol (HFIP; MilliporeSigma; cat. no. 105228) to a final concentration of 1 mg/mL. The HFIP/peptide mixture was incubated at room temperature for 1 h and sonicated for 10 min in a water-bath sonicator. The mixture was then air-dried under a gentle stream of nitrogen gas for 1.5 h. The peptide crystal was re-suspended in 100 % DMSO, aliquoted to 10 µL per tube and stored at –80 °C until experimental use (see next section).

### Hippocampal slice preparation

Mice were deeply anaesthetized via inhalation with a mixture of 5 % isofluorane / 95 % oxygen and euthanized by cervical dislocation. The brain was rapidly removed (<1 min) and chilled in cold ACSF solution (containing, in mM: 124 NaCl, 3 KCl, 26 NaHCO_3_, 1.25 NaH_2_PO_4_, 10 D-glucose, 2 MgSO_4_ and 2 CaCl_2_; bubbled continuously with 95% O_2_ / 5% CO_2_). Hippocampi were isolated from both hemispheres and transverse slices (400 µm thickness) were prepared from the dorsal ends of both hippocampi using a Mcllwain tissue chopper (Ted Pella; Redding, California). The CA3 region was left intact for LTP but removed via a surgical cut in LTD experiments. Slices were transferred to a holding chamber with room temperature ACSF and allowed to recover for at least 1 h. On the day of experiment, the Aβ peptide/DMSO mixture was diluted to 100 µM in D-PBS, vortexed briefly and allowed to aggregate for 2 h at room temperature to form oAβ. The oAβ stock was then diluted to a final concentration of 500 nM in oxygenated ACSF and acutely applied to hippocampal slices for 2 h at room temperature. As an inclusion criterion to account for batch-to-batch variability, we routinely assessed the bioactivity of our oAβ preparation by confirming the ability of each oAβ batch to impair hippocampal LTP (Figure 1A,B).

**Figure 1:**
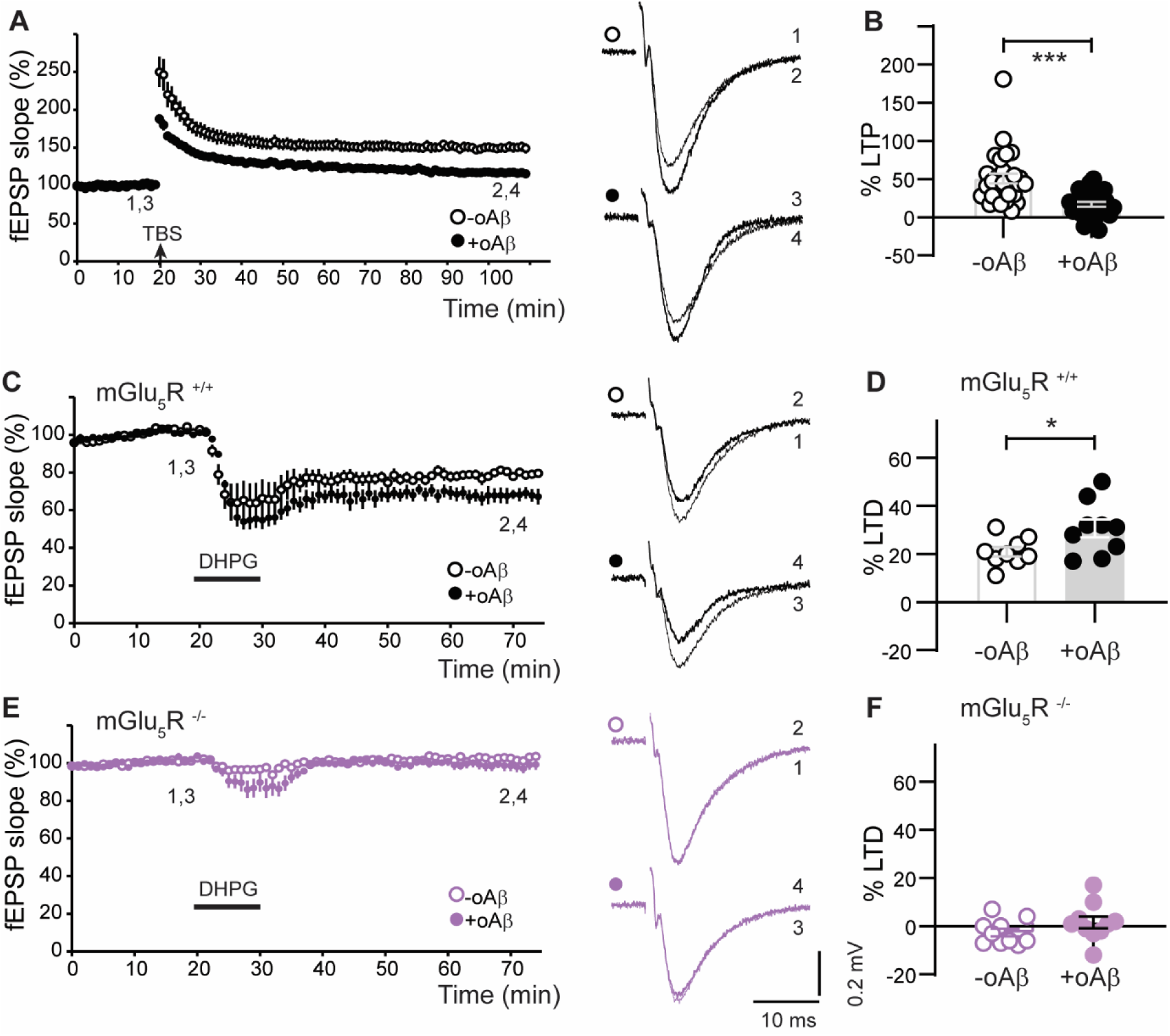
oAβ enhances mGlu_5_R-LTD at CA1 synapses. (**A-B**) Validation of oAβ preparation, confirming its ability to impair LTP. Delivery of a theta-burst protocol (TBS; see Methods), at the time indicated by the arrow, induced LTP in –oAβ slices (51 ± 7 %, n = 28). LTP was impaired in the interleaved +oAβ slices (17 ± 3 %, n = 28). Representative fEPSP traces for both conditions are shown to the right of the time course plot (superimposed for sample experiment, before and after TBS, at times indicated by the numbers 1-4). **B** Bar graph shows the level of LTP in each experiment. p = 0.00002. (**C-D**) Application of DHPG (50 µM, 10 min), induced LTD in –oAβ slices from mGlu_5_R^+/+^ mice (21 ± 2 %, n = 9). Interleaved +oAβ slices showed an enhancement of DHPG-LTD (31 ± 4 %, n= 9) with summary of LTD experiments in **D**. p = 0.032. Representative fEPSP traces demonstrating enhanced DHPG-LTD in +oAβ compared to –oAβ slices. (**E-F**) In slices from littermate mGlu_5_R^-/-^ mice, DHPG failed to induce LTD in either condition (+oAβ = 2 ± 3 %, n=10; –oAβ = –3 ± 2 %, n=10) with summary in **F**. Representative fEPSP traces showing the absence of DHPG-LTD in both conditions. Sample fEPSP traces in all panels are the mean of 4 consecutive responses at the indicated time points (1-4). For each experiment, sample traces before and after DHPG treatment are superimposed. Two-tailed t-tests were used in **B**, **D** and **F**.

### Electrophysiology

After vehicle or oAβ incubation, a dorsal hippocampal slice was submerged in a Slicemate recording chamber (Scientifica; Uckfield, U.K.) with ACSF perfusing at a rate of 2.5 mL/min and temperature maintained at 30.0 ± 0.5 °C. Extracellular field potentials were recorded from the stratum radiatum area of CA1 using a glass microelectrode (∼2-3 MΩ) filled with ACSF. Responses were evoked by stimulating the Schaffer collaterals using a platinum-iridium bipolar electrode (FHC; Bowdoin, Maine) positioned at the border between area CA2/CA1. Stimuli of 100 µs in duration were delivered once every 15 s via a DS2 constant voltage isolator (Digitimer; Hertfordshire, U.K.). Responses were amplified using a Multiclamp 700B amplifier (Molecular Devices; San Jose, California), digitized at 40 kHz and monitored in real time using WinLTP software (WinLTP Ltd.: Bristol, U.K: Anderson & Collingridge, 2007). After a period of stable baseline recordings, datapoints for input/output (I/O) plots were obtained with fixed stimulus intensity at 1x, 1.5x, 2x, 3x, and 4.5x the threshold for evoking a visually detectable field excitatory postsynaptic potential (fEPSP) (≤ 0.05 mV). For plasticity experiments a stimulus intensity of 3x the threshold intensity was employed. LTP was induced by a theta-burst stimulation (TBS) protocol, with four stimuli delivered at 100 Hz, repeated 10 times at a frequency of 5 Hz. LTD was chemically induced by bath-application of (S)-3,5-DHPG (50 µM; Abcam; Cambridge, U.K.; cat. no. ab120007) for 10 min. For offline analysis, responses were low-pass filtered at 2 kHz and the mean of four consecutive responses was quantified.

### Quantification and statistical analysis

Data in the time course plots were normalized to the mean of the entire baseline period (defined as 100 %) and presented as mean ± standard error of mean (SEM). The magnitude of LTP or LTD was quantified and presented as the percentage change of the fEPSP slope within the last 5 min of the recording *versus* the baseline period. Fiber volleys (FVs) and fEPSPs were quantified using peak amplitude and initial slope measurements, respectively, using WinLTP. The I/O plots were analysed using Microsoft Excel linear regression software. The respective slope values from a fitted line between FV *versus* stimulation intensity and fEPSP slope *versus* FV were used for quantification. Statistical comparisons between –oAβ and +oAβ conditions were assessed using an unpaired two-tailed Student’s t-test. The level of significance was set at p<0.05.

## Results

### Acute oAβ exposure enhances mGlu_5_R-dependent LTD

To investigate the impact of oAβ on synaptic function, we incubated acute hippocampus slices for 2 h with oAβ (500 nM). We subsequently measured synaptic plasticity with field potential recordings at Schaffer collateral-CA1 synapses. As a positive control for the bioactivity of our oAβ preparation, we confirmed previous findings (Shankar et al., 2008; Li et al., 2011) that oAβ was able to impair LTP in slices pretreated with oAβ (+oAβ) *versus* interleaved non-treated slices (-oAβ; Figure 1A,B). Although LTP was impaired by acute application of oAβ, basal synaptic transmission was unaltered, indicated by the overlapping input-output relationship of +oAβ compared to –oAβ slices (Figure S1). As such, this paradigm allowed us to study early disruptions to plasticity that precedes the loss of synapses.

To investigate how oAβ alters mGluR-dependent synaptic plasticity mechanisms, we induced LTD using DHPG, a selective agonist for group I mGluRs (mGlu_1_R and mGlu_5_R), that is widely used to study mGluR function. A short application of DHPG (10 min) induced a stable LTD (DHPG-LTD) in –oAβ slices from wild-type mice (Figure 1C,D). The magnitude of DHPG-LTD was significantly enhanced in the interleaved +oAβ slices (Figure 1C,D), consistent with previous observations (Hsieh et al., 2006; Shankar et al., 2008; Li et al., 2009; Cavallucci et al., 2013; Mango & Nisticò, 2018; Yang et al., 2021). To determine the group I mGluR subtype-specificity of our observed phenotype, we used hippocampal slices from mGlu_5_R^-/-^ mice (Lu et al., 1997) which were littermates to the mGlu_5_R^+/+^ mice in Figure 1C,D. DHPG-LTD was absent in both –oAβ and +oAβ slices (Figure 1E,F). This indicates that the form of LTD which is enhanced by oAβ requires the activation of mGlu_5_ receptors (referred to hereafter as mGlu_5_R-LTD). That is, both the LTD under control conditions and the additional LTD observed in the presence of oAβ are both fully dependent on the activation of mGlu_5_Rs.

### NMDARs are required for oAβ-mediated enhancement of mGlu_5_R-LTD

There are two distinct forms of DHPG-LTD that can be distinguished by their dependence upon the activation of NMDARs (Palmer et al., 1997; Sanderson et al., 2022). To determine whether NMDARs are required for mGlu_5_R-LTD either under control conditions or the enhanced mGlu_5_R-LTD observed in the presence of oAβ, we used a selective NMDAR antagonist, L689,560 (5 µM). L689,560 was used as it acts at the glycine site of the NMDAR and so is independent of L-glutamate concentration, which might be altered by oAβ treatment (Li et al., 2009). In these experiments, we interleaved a new set of controls and again observed the potentiation of mGlu_5_R-LTD by oAβ (Figure 2A,B). We found that NMDAR activation was not required for mGlu_5_R-LTD in control (-oAβ) conditions. However, the enhancement of mGlu_5_R-LTD by oAβ was completely prevented by L689,560 (Figure 2C,D) and therefore requires NMDAR signaling.

**Figure 2:**
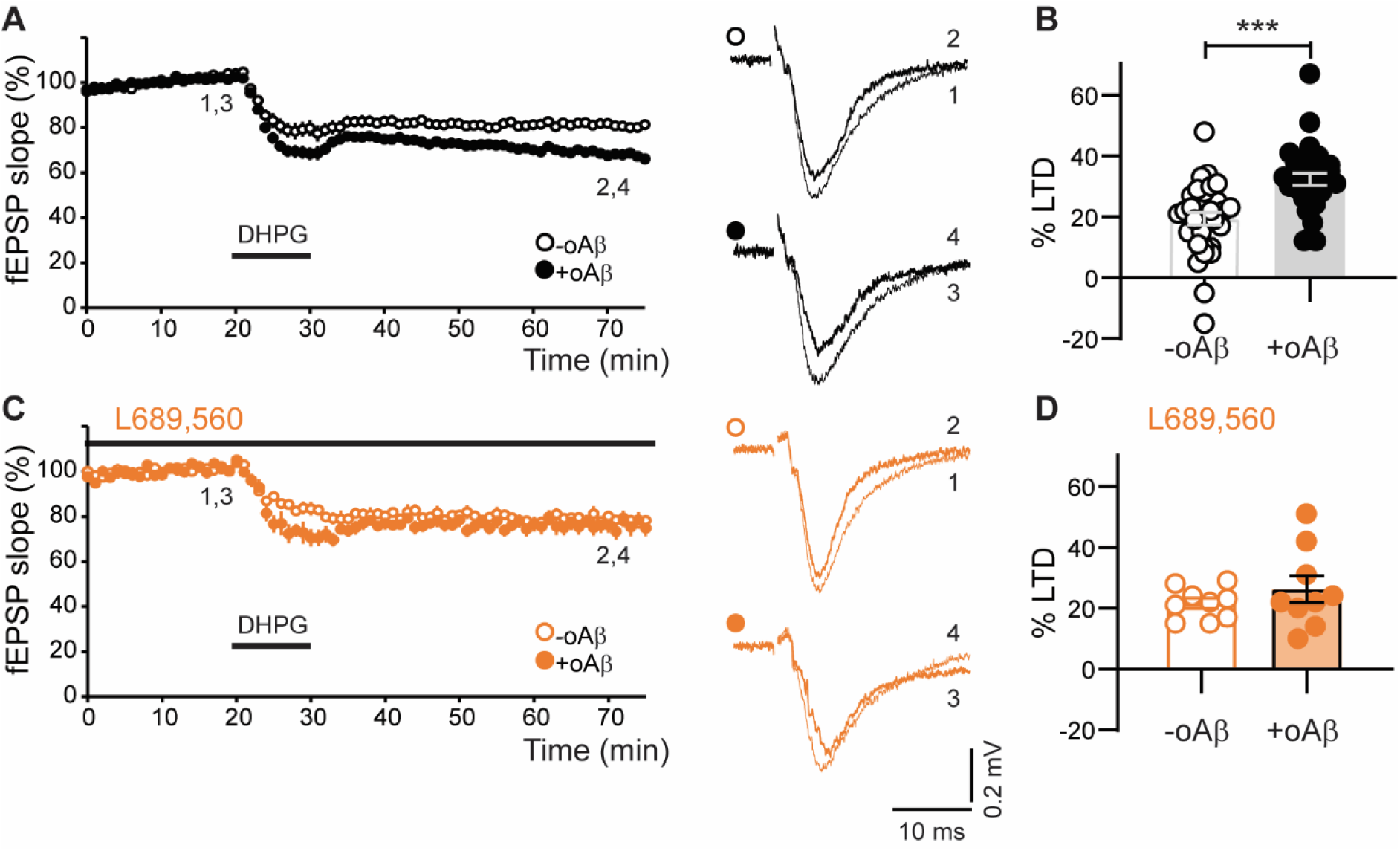
Enhancement of mGlu_5_R-LTD by oAβ requires NMDARs. (**A-B**) In a separate cohort, pretreatment with oAβ also resulted in an enhancement of DHPG-LTD (34 ± 2 %, n=22) compared to the interleaved –oAβ slices (21 ± 2 %, n=22). p=0.00031. (**C-D**) In the presence of L689,560 (5 µM), the magnitude of DHPG-LTD was similar in +oAβ slices (26 ± 4 %, n=9) and –oAβ slices (22 ± 2 %, n=9). Sample fEPSP traces in all panels are the mean of 4 consecutive responses at the indicated time points (1-4). For each experiment, sample traces before and after DHPG treatment are superimposed. A two-tailed t-test was used in **B** and **D**.

### oAβ enhances mGlu_5_R-LTD via complement cascade signaling

Early studies of the effects of oAβ on synaptic plasticity found that inhibition of LTP was prevented by application of minocycline, an inhibitor of microglia activation (Wang, Rowan, & Anwyl, 2004). Further, it has recently been found that agonism of group I mGluRs *in vivo* activates the complement cascade leading to synapse loss (Bie et al., 2019). We therefore hypothesized that complement signaling is necessary for the enhancement of mGlu_5_R-LTD by oAβ. We antagonized either C3aR or C5aR, as both receptors have been implicated in the pathology of genetic mouse models of AD (Fonseca et al., 2009; Lian et al., 2016; Hernandez et al., 2017; Litvinchuk et al., 2018; Carvalho et al., 2022; Gomez-Arboledas et al., 2022). For these experiments we also interleaved a new set of controls (Figure 3A,B) and again observed the potentiation of mGlu_5_R-LTD by oAβ. To inhibit C5aR, we bath-applied PMX205 (0.3 µM), a peptide-based antagonist (March et al., 2004). PMX205 had no effect on the level of mGlu_5_R-LTD induced under control conditions but eliminated the enhancement of LTD induced by oAβ (Figure 3C,D). We next used W54011, a non-peptide based C5aR antagonist (Sumichika et al., 2002). As with PMX205, bath-application of W54011 (0.3 µM) had no effect on the level of mGlu_5_R-LTD induced under control conditions but eliminated the potentiation induced by oAβ (Figure 3E,F). Lastly, we investigated the role of C3aR using the antagonist SB290157 (Ames et al., 2001). As with the C5aR antagonists, SB290157 (1 µM) abolished the enhanced mGlu_5_R-LTD induced by oAβ without affecting the level of mGlu_5_R-LTD under control conditions (Figure 3G,H). Together, these data indicate that the oAβ-mediated enhancement of mGlu_5_R-LTD occurs via activation of complement C3aR/C5aR signaling.

**Figure 3:**
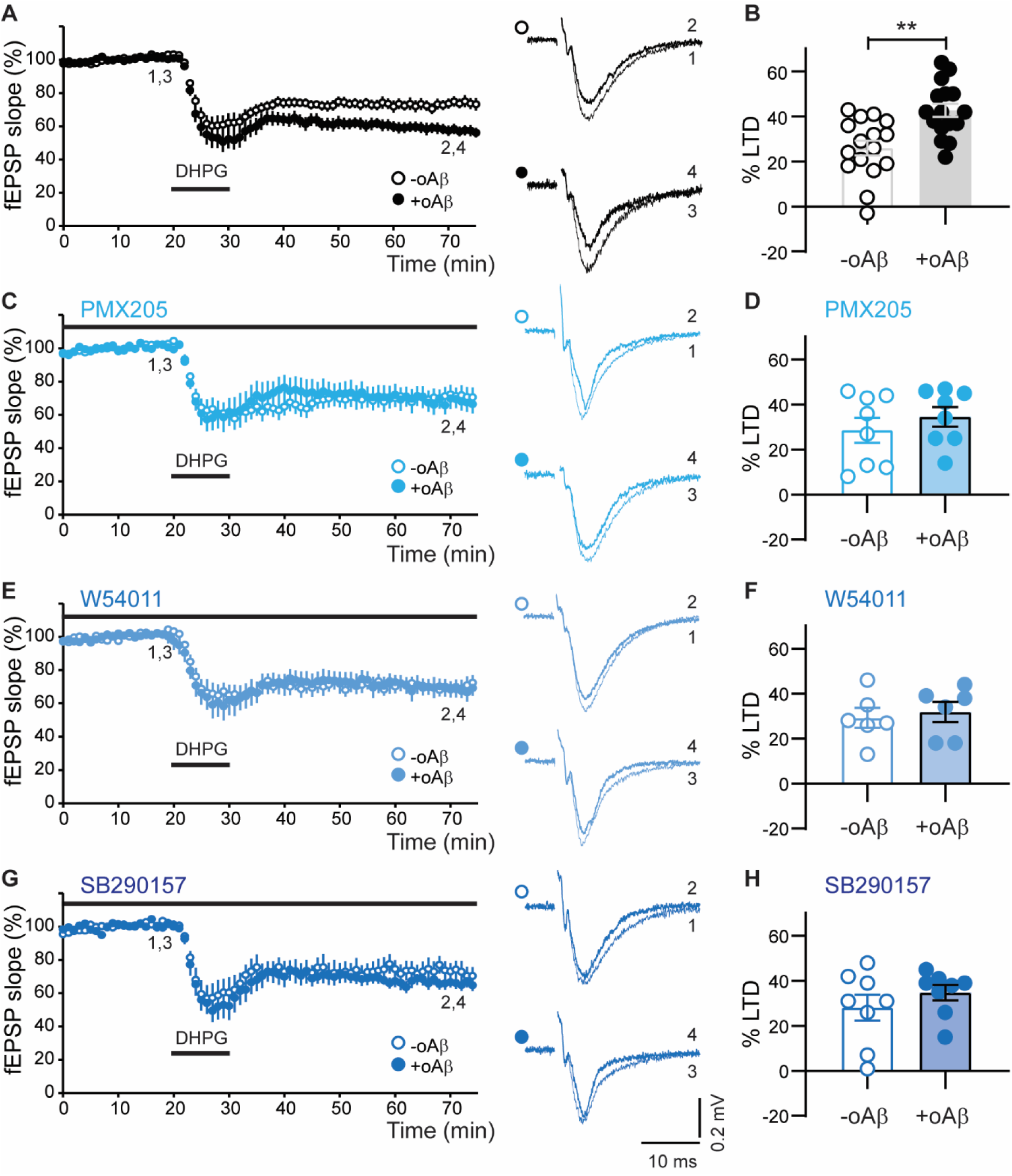
Complement C5aR and C3aR signaling is necessary for oAβ to enhance mGlu_5_R-LTD. (**A-B**) In a third cohort, pretreatment with oAβ enhanced DHPG-LTD (43 ± 3 %, n=16) compared to the interleaved –oAβ slices (26 ± 3 %, n=16). p=0.00064. (**C-D**) In the presence of the C5aR antagonist PMX205 (0.3 µM), the level of DHPG-LTD was similar in +oAβ slices (35 ± 4 %, n=8) and –oAβ slices (29 ± 6 %, n=8). (**E-F**) In the presence of a separate C5aR antagonist, W54011 (0.3 µM) the magnitude of DHPG-LTD was similar in +oAβ slices (32 ± 5 %, n=6) compared to –oAβ slices (29 ± 4 %, n=6). (**G-H**) In the presence of a C3aR antagonist, SB290157 (1 µM), DHPG-LTD magnitude was not significantly different between +oAβ slices (35 ± 3 %, n=8) and –oAβ slices (28 ± 6 %, n=8). Sample fEPSP traces in all panels are the mean of 4 consecutive responses at the indicated time points. For each experiment, sample traces before and after DHPG treatment are superimposed. Two-tailed t-tests were used in **B**, **D**, **F** and **H**.

## Discussion

The complement cascade is emerging as a key innate immune pathway for shaping synapses during brain development, as well as driving synapse loss in various disease states. However, the synaptic signals that lead to complement activation remain poorly understood. In the present study, we uncover a mechanistic link between innate immune signaling and the amplification of glutamatergic synapse depression induced by oAβ. We found that mGlu_5_R-dependent LTD induced by DHPG application was enhanced in slices treated with oAβ (Figure 1). Unlike vehicle-treated slices, the oAβ-mediated enhanced portion of mGlu_5_R-LTD was dependent on NMDAR signaling (Figure 2). Similarly, antagonism of either C3aR or C5aR prevented oAβ from enhancing mGlu_5_R-LTD (Figure 3), without affecting the underlying mGlu_5_R-LTD *per se*. Given that basal synaptic transmission was unaltered by oAβ (Figure S1) our findings reveal a signaling axis between mGluRs, NMDARs, and the complement cascade which mediates early synaptic plasticity alterations, upstream of overt synapse loss.

### oAβ-induced glutamatergic synapse dysfunction

Although AD is an extremely slow progressing disease, it is likely that at the level of a single synapse there is acute damage, which in many cases may be orchestrated by toxic oAβ species. This would presumably occur when the processes that ordinarily prevent the accumulation of oAβ are sufficiently compromised such that the local concentration increases to a level that can impair synaptic function and structure. Therefore, the transient application of oAβ has been widely employed to model this early, critical stage of the disease. Consistent with the validity of this model, mechanisms that are engaged by the acute application of oAβ to impact synaptic plasticity, such as the involvement of GSK-3, caspase-3 (Jo et al., 2011), tau (Shipton et al., 2011; Taylor et al., 2021) and microglia (Wang et al., 2004) are all directly relevant to human AD pathology.

Multiple studies have found that oAβ causes a shift in the LTP-LTD balance, favoring LTD (Cullen, Suh, Anwyl, & Rowan, 1997; Walsh et al., 2002; Hsieh et al., 2006; Shankar et al., 2007; Shankar et al., 2008; Li et al., 2009; Jo et al., 2011; Li et al., 2011; Cavallucci et al., 2013). Altered activity of both group I mGluRs and NMDARs by oAβ is central to this glutamatergic synaptic failure. mGlu_5_R was identified as a co-receptor of cellular prion protein (PrP_c_) for binding oAβ and is necessary for oAβ-induced intracellular signal transduction and synapse loss (Um et al., 2013; Haas, Kostylev, & Strittmatter, 2014; Beraldo et al., 2016; Brody & Strittmatter, 2018). Additionally, oAβ binds to NMDARs (De Felice et al., 2007; Decker et al., 2010) and impairs NMDAR-dependent LTP while enhancing NMDAR-dependent LTD (Cullen et al., 1997; Walsh et al., 2002; Wang et al., 2004; Hsieh et al., 2006; Shankar et al., 2007; Shankar et al., 2008; Jo et al., 2011; O’Riordan, Hu, & Rowan, 2018; Abd-Elrahman et al., 2020; Taylor et al., 2021). Despite these important advances, whether NMDARs and mGluRs are synergistically involved in oAβ-induced synaptic weakening and loss is poorly understood. Of relevance to the present study, at least two distinct forms of DHPG-LTD exist, which can be distinguished by the dependence on NMDARs (Palmer et al., 1997; Sanderson et al., 2022). In our current study, DHPG-LTD induction under control conditions was not affected by NMDAR antagonism. In contrast the oAβ-enhanced portion of the LTD fully depended on the activation of NMDARs. As such, promotion of mGluR-NMDAR signaling during synaptic depression may be a key aspect of the shift in LTP/LTD balance caused by oAβ.

Both mGlu_5_R and NMDARs are also implicated in AD pathogenesis in genetic mouse models and humans. Antagonists of mGlu_5_R and genetic deletion have both been found to reduce synaptic and behavioral deficits in AD mouse models (Hamilton et al., 2014; Hamilton et al., 2016; Haas et al., 2017; Abd-Elrahman et al., 2020; Spurrier et al., 2022). Indeed, the mGlu_5_R SAM BMS-984923 (Haas et al., 2017), is currently in phase I clinical trials. Evidence that altered NMDAR signaling is causally related to the human condition is the use of memantine, a non-competitive NMDAR antagonist, for the treatment of AD. Memantine slows the progression of dementia and its underlying therapeutic mechanism has been attributed to the normalization of synaptic plasticity (Parsons, Stöffler, & Danysz, 2007; Wang & Reddy, 2017; Liu, Chang, Song, Li, & Wu, 2019). Our experiments reveal one way in which NMDARs and mGlu_5_Rs may act synergistically to control synaptic integrity.

### C3aR/C5aR signaling in AD

In both post-mortem human samples as well as numerous AD mouse models, many components of the complement cascade have been found to be upregulated (Hammond et al., 2018; Schartz & Tenner, 2020). Importantly, pharmacological and genetic manipulations of C3aR and C5aR have provided evidence that both receptors are necessary for synaptic dysfunction and cognitive impairments in AD genetic mouse models. In particular, the C3aR and C5aR antagonists used in the current study, SB290157 and PMX205, respectively, have each been found to reduce pathology in genetic mouse models of AD (Fonseca et al., 2009; Lian et al., 2016; Gomez-Arboledas et al., 2022). Furthermore, synapse engulfment by microglia and neuron loss are reduced in C3aR^-/-^ mice in the PS19 tauopathy AD model (Litvinchuk et al., 2018) and C5aR1 genetic ablation is protective in AD mouse models (Hernandez et al., 2017; Carvalho et al., 2022). However, the role of oAβ in aberrant C3aR/C5aR signaling, and the intersection with glutamatergic synaptic signaling to cause this dysfunction was previously not understood. Our study provides insight into this communication, by demonstrating that oAβ enhances synaptic weakening, via NMDAR and mGlu_5_R synergistic mechanisms that require both C3aRs and C5aRs.

One important question to be addressed by future studies is which arm(s) of the complement cascade initiates the cleavage of C3 following mGlu_5_R-LTD in the presence of oAβ? Proteolytic cleavage of C3 generates both C3a, as well as components of the C5 convertase which produces C5a. C3 conversion can be driven by either the classical arm, initiated by C1q, the lectin arm via mannose-binding lectin (MBL) or the alternative pathway through C3 “tickover” (Ricklin et al., 2010). Hong et al. (2016) found that the ability of oAβ to inhibit LTP requires the classical complement pathway, as co-incubation of hippocampal slices with oAβ and a function-blocking antibody of C1q restored LTP. Furthermore, *in vivo* injections of oAβ into rat hippocampus upregulated C1q, and increased synapse phagocytosis by microglia (Bie et al., 2019). C1q upregulation by oAβ and subsequent synapse phagocytosis required group I mGluR signaling, and *in vivo* DHPG administration alone increased C1q. Finally, a recent study found that a SAM of mGlu_5_R reduced C1q co-localization with PSD-95 and microglia engulfment of synapses in a mouse model of AD (Spurrier et al., 2022). As such, our findings suggest that C3aR/C5aR activation links the induction of the classical arm of the complement cascade, via mGluRs and NMDARs, to downstream synaptic depression.

Cleavage of C3 also generates the C3b fragment, which can covalently attach to synapses where it is converted to iC3b and binds to the phagocytic receptor CR3 (Ricklin et al., 2010). Genetic deletion of CR3 has also been found to spare synapses following oAβ administration *in vivo* (Hong et al., 2016), indicating that phagocytosis through iC3b-CR3 binding is also critical for the deleterious effects of complement in pathology. As weaker synapses are targeted by complement for phagocytic removal (Schafer et al., 2012), an intriguing question is whether synapse weakening by C3aR/C5aR activation also tags those synapses with iC3b for subsequent engulfment by microglia. CR3 can also impact synapses by non-phagocytic mechanisms in neuroinflammatory conditions. LPS and hypoxia activates CR3 which induces LTD of AMPAR transmission in the hippocampus (Zhang et al., 2014). Thus, microglia and the complement cascade have a multi-faceted role in synaptic dysfunction in disease, being critical for both changes to AMPAR-mediated transmission strength as well as structural synapse elimination.

### Relevance to other neurodegenerative diseases

Upregulation of complement cascade proteins and activity is not exclusive to AD and has been detected in several other neurodegenerative diseases. For example, A1 reactive astrocytes, which express high levels of C3, have been identified in human post-mortem tissue from patients with amyotrophic lateral sclerosis, Huntington’s disease, Parkinson’s disease, and multiple sclerosis (Liddelow et al., 2017). Further, studies from *in vitro* and mouse models of these diseases have demonstrated the necessity of C3 and/or C5 for their pathology (Woodruff et al., 2008; McGeer, Lee, & McGeer, 2017; Wang, Lee, Lee, Woodruff, & Noakes, 2017; Hammond et al., 2020; Bourel et al., 2021; Chen, Hu, Ding, Du, & Hu, 2021; Gregersen et al., 2021; Lopez-Sanchez et al., 2022). Interestingly, mGlu_5_R has separately been identified as an emerging therapeutic target in these complement-associated neurodegenerative diseases (Geurts et al., 2003; Abd-Elrahman et al., 2017; Bonifacino et al., 2017; Doria et al., 2018; Bonifacino et al., 2019; Farmer et al., 2020; Zhang et al., 2021; Azam et al., 2022; Li, Colson, Abd-Elrahman, & Ferguson, 2022; Su, Wang, Han, & Shen, 2022). Therefore, the pathological mGlu_5_R-NMDAR-C3aR/C5aR signaling axis identified in our study may have relevance for multiple brain disorders.

### Concluding Remarks

mGluRs, NMDARs, and the complement cascade have been independently studied for their role in mediating the deleterious effects of oAβ. Our study has found a direct link which places glutamate receptor-innate immune interactions at the center of early synaptic dysfunction. As oAβ is thought to be a key trigger in AD, a deeper understanding of the cellular and molecular mechanisms of these interactions has the potential to elucidate therapeutic targets which halt AD progression.

## Acknowledgements

This work was supported by a CIHR (Canadian Institutes of Health Research) Foundation Grant to GLC (#154276). GLC is the holder of the Krembil Family Chair in Alzheimer’s Research. We thank the trainee support provided by the C.R. Younger Foundation. The work was supported by the MS Society of Canada as well as gifts from Jon and Nancy Love and the Dani Reiss Family Foundation (Neurodegeneration and Aging Research Program). We are grateful for the long-standing support of the Amalgamated Transit Union (ATU) Local 113 (Toronto, Canada). EWS was supported by a CIHR Doctoral Award (Frederick Banting and Charles Best Canada Graduate Scholarship, CGS-D) and an Ontario Graduate Scholarship (OGS).

---

**Supplemental Figure 1:**
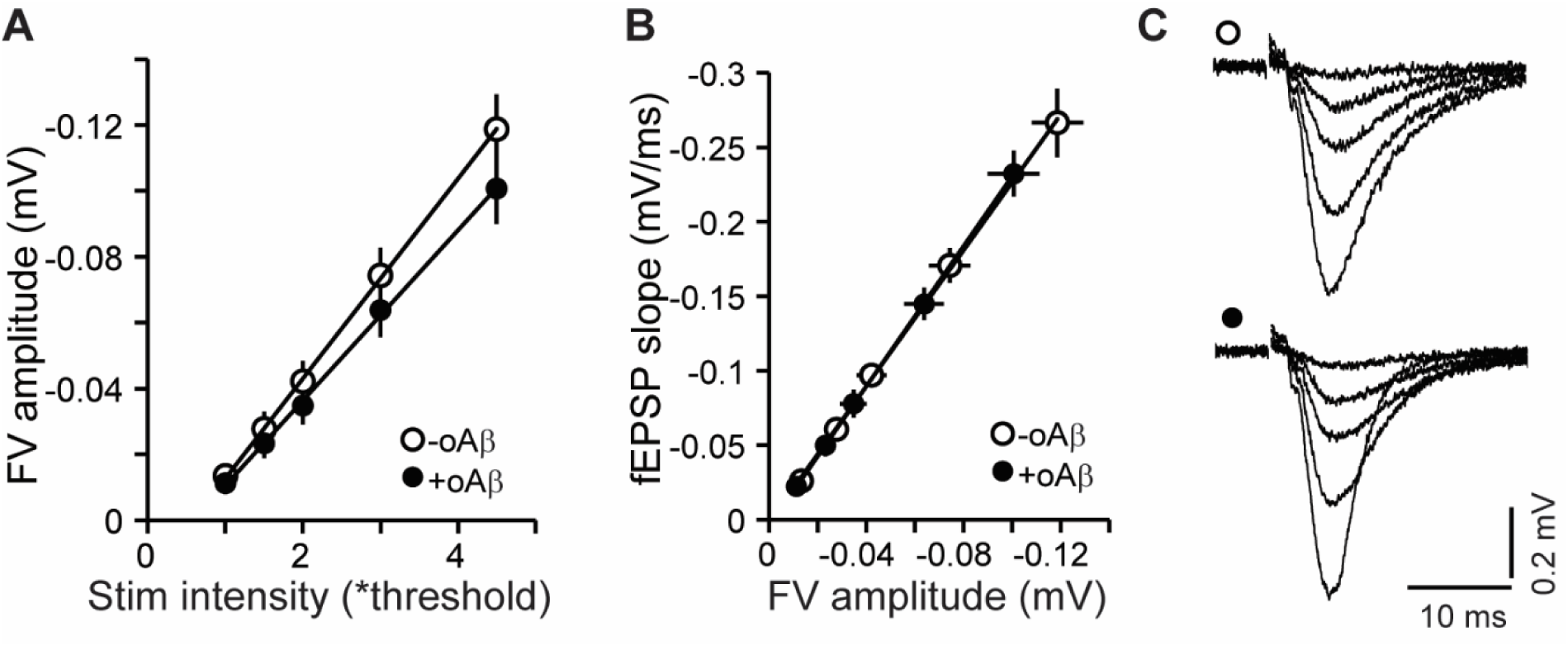
Acute oAβ application does not affect basal synaptic strength at CA1 synapses. (**A-C**) CA3-CA1 basal synaptic transmission is similar in +oAβ slices and –oAβ slices, as assessed by input-output curves. **A** Fiber volley (FV) amplitude *versus* stimulus intensity (line slopes: +oAβ = –0.026 ± 0.003, n = 13; –oAβ = –0.030 ± 0.003, n = 13). **B** fEPSP slope *versus* FV amplitude (line slopes: +oAβ = 2.517 ± 0.237, n = 13; black, *versus* –oAβ = 2.513 ± 0.362, n = 13; white). **C** Representative fEPSP traces at different stimulus intensity (multiples of threshold) for both conditions.

